# A lightweight, physics-based, sensor-fusion filter for real-time EEG denoising and improved downstream AI classification

**DOI:** 10.1101/2025.09.24.675953

**Authors:** Jack-Michael Wesierski, Nicholas Rodriguez

## Abstract

Physiological time-series data, like electroencephalography (EEG), are vulnerable to motion, ocular, and muscle artifacts that hinder real-time inference and bias offline analyses. We present the Minds AI Filter: a lightweight, physics-based, sensor-fusion method that exploits multichannel spatial structure and band-aware synchrony to enhance neural activity while suppressing non-neural noise. The nomenclature “AI” reflects integration within a larger artificial-intelligence pipeline; the filter itself requires no prior training or deep learning. A single tuning parameter controls filter strength. The design supports streaming windows (*≈*1 s) with minimal added latency and extends naturally to longer offline segments; leveraging a sensor-fusion design across channels, it suggests applicability to other neurophysiological time-series, such as MEG and ECoG, pending further validation; exploratory incorporation of EOG/ECG as auxiliary signals is a potential avenue for future filter advancements.

We evaluate the approach across multiple devices and public datasets, assessing both down-stream AI classification performance and real-time signal-quality metrics. In both real-time and offline settings, the filter performed better on dynamic artifacts and noise than baseline and commonly used alternatives in our evaluations. When applied in conjunction with other methods, it was only observed to improve downstream accuracy, never reduce it, when any effect was present. Denoising is quantified using SNR-like measures, and ablations isolate the roles of spatial coupling and band weighting. Artifact-specific analyses (ocular bursts, head tilt, jaw clench) and latency profiling on commodity hardware are included. These results indicate that a lightweight, synchrony-aware filter can robustly stabilize real-time EEG and systematically improve downstream AI classification. The method is compatible with standard preprocessing but does not depend on it.

## 1 Introduction

Electroencephalography (EEG) enables non-invasive access to brain dynamics for applications such as affect recognition, inner-speech interfaces, and motor-imagery control, but its low amplitude and non-stationarity make it highly susceptible to motion, ocular, and myogenic artifacts, especially in real-world, real-time use. Conventional preprocessing (e.g., notch/band-pass, common average referencing, PCA/ICA, adaptive filters, ASR) can be effective yet often involves assumptions or supervision (clean baselines, component labeling) that are difficult to satisfy online [2–5, 8].

We introduce the Minds AI Filter: a physics-informed, multichannel denoising method that leverages synchrony across channels to suppress non-neural components while preserving neural structure [1, 7, 9, 10]. “AI” is included in the naming convention due to its part in a larger artificial intelligence pipeline, but the filter does not require prior training or deep learning. The filter is controlled by a single scalar hyperparameter (*λ*), runs efficiently on short streaming windows, and extends naturally to offline analysis. It can be used on its own and remains compatible with standard preprocessing steps. Although developed for EEG, the design may extend to MEG and ECoG, which share multichannel oscillatory structure; we also anticipate exploratory incorporation of EOG/ECG as auxiliary signals in future filter advancements. Formal validation is warranted.

### Contributions

(i) a multichannel, synchrony-aware filtering approach; (ii) a simple method with a single hyperparameter (*λ*) suitable for streaming; (iii) comprehensive evaluation across devices, datasets, and downstream AI classification; and (iv) ablation and latency measurements demonstrating real-time viability.

## 2 Related Work

Adaptive filtering (e.g., LMS/RLS), subspace methods (PCA/ICA), spatial filters (CAR/Laplacian), and artifact subspace reconstruction (ASR) are widely used [2–5, 8]. Synchrony-aware and physics-informed formulations, including oscillator-coupling perspectives related to the Kuramoto family, offer complementary priors for neural denoising and structure discovery [1, 7, 9, 10]. Our approach differs by providing a single-hyperparameter, streaming-friendly filter that composes cleanly with downstream pipelines and artifact-specific steps, without requiring component labeling.

## 3 Methods (high-level)

### 3.1 Signal model and objective

Let *X ∈* ℝ^*C×T*^ denote multi-channel EEG. We estimate a denoised 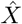 by minimizing an objective combining (i) a data-fidelity term with (ii) a regularizer that promotes physiologically plausible synchrony across channels. The regularizer has two parts: (a) **spatial coupling** informed by electrode topology or learned correlations, and (b) **band-aware** weighting to reflect distinct spatial scales in canonical frequency bands. A single scalar *λ* controls overall regularization strength. A minimal reference implementation is provided in **Supplementary File S1** (`filter.csv`).

### 3.2 Connectivity prior & band weighting

Electrode geometry (10–20/10–10) or empirical correlations define a normalized connectivity prior. We construct a *k*-nearest-neighbor channel graph (*k ≈* 4–6) with distance-decayed weights and use the normalized Laplacian to encourage band-limited spatial compatibility among neighboring channels. Band weights (*w*_*b*_) reflect physiology—stronger coupling for delta/theta, moderate for alpha/beta, lighter for gamma—and node-degree normalization provides boundary correction so edge electrodes are not over-regularized.

### 3.3 Hyperparameter (*λ*) and streaming

*λ* modulates aggressiveness vs. preservation. For low-latency windows (*≈*1 s), tighter settings are effective; longer windows (*≥*30–60 s) typically favor more conservative settings. In practice, tuning with window length and sampling rate is recommended. Overlapped streaming windows maintain continuity; no component labeling is required.

### 3.4 Evaluation metrics

#### Offline Artificial Intelligence

- Balanced accuracy of emotional valence classification using logistic regression on features derived from wavelet coefficients. Further technical architecture will be included in the final submission.

#### Real-time signal-quality (per *∼*1 s window unless noted)

- **SNR (power ratio):** 𝔼[*s*^2^]*/*𝔼[*n*^2^] (optional dB = 10 log10(·)); where *s* = *x*_filt_ and *n* = *x*_raw_ *− x*_filt_.
- **SNR (amplitude ratio):** 𝔼[|*s*|]*/*𝔼 [|*n*|] (optional dB = 20 log_10_(·)).
- **SNR (variance ratio):** Var(*s*)*/*Var(*n*) (optional dB = 10 log_10_(·)).
- **Artifact suppression (peak drop):** 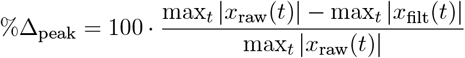; default flag at *≥* 20% reduction.
- **Baseline drift (mean/median shift):** Δ_*µ*_ = *µ*_filt_ *− µ*_raw_, Δ_med_ = med_filt_ *−* med_raw_; default flag at |shift| *≥* 5 *µ*V.
- **Variance smoothing:** 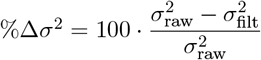; default flag at *≥* 5% reduction.
- **Channel aggregation:** metrics may be computed per channel and/or on the mean across EEG channels only.

Implementation details for these streaming metrics are provided in **Supplementary File S2** (`Minds_AI_Filter_Real-time_Signal_Analysis.txt`).

### Reference implementation

We include a reference Python implementation of the Minds AI Filter as **Supplementary File S1** (`filter.txt`). The public method is a single convenience function that operates directly on a 2-D array of shape **[channels, timepoints]**:

~~~
import mindsai_filter_python as mai
# X: numpy array shaped [channels, timepoints]
X_filt = mai.mindsai_python_filter(X, lambda_val=1e-25)
~~~

The code computes channel phases via the analytic signal (Hilbert transform), constructs a synchrony operator from phase differences, and applies a closed-form multichannel filter controlled by *λ* (see Listing S1 for details). This function is **stateless** and requires no model fitting or prior training. The supplemental file is versioned with the preprint; a persistent archive (DOI) will be added upon submission.

## 4 Datasets & Experimental Setup

### Devices

OpenBCI Cyton+Daisy (16 channels, 125 Hz), Neurosity Crown (8 channels, *≈*256 Hz), BrainBit Dragon (*≥*16 channels, up to 500 Hz).

### Public datasets

Affective state datasets such as DEAP (32 subjects, 32 channels & 512 Hz) [6], and others where indicated.

### Paradigms & artifacts

Resting/task segments; eyebrow raises, head tilt/motion, blinks, jaw clench.

### Baselines

Raw, notch+band-pass, CAR/Laplacian, PCA/ICA (with standard component rejection), ASR, LMS-style adaptive filters.

### Ablations

(i) remove connectivity prior; (ii) use uniform band weights; (iii) sweep *λ*.

## Results

### 5.1 Real-time denoising & signal quality

On short windows (*≈*1 s), the filter suppresses motion- and ocular-locked bursts while preserving underlying rhythms. SNR-like metrics (power, amplitude, variance ratios), baseline-drift reduction, and variance smoothing improve consistently across devices. Representative traces (raw, filtered, residual) and per-channel summaries are provided in figures; see Fig. 1.

**Figure 1:**
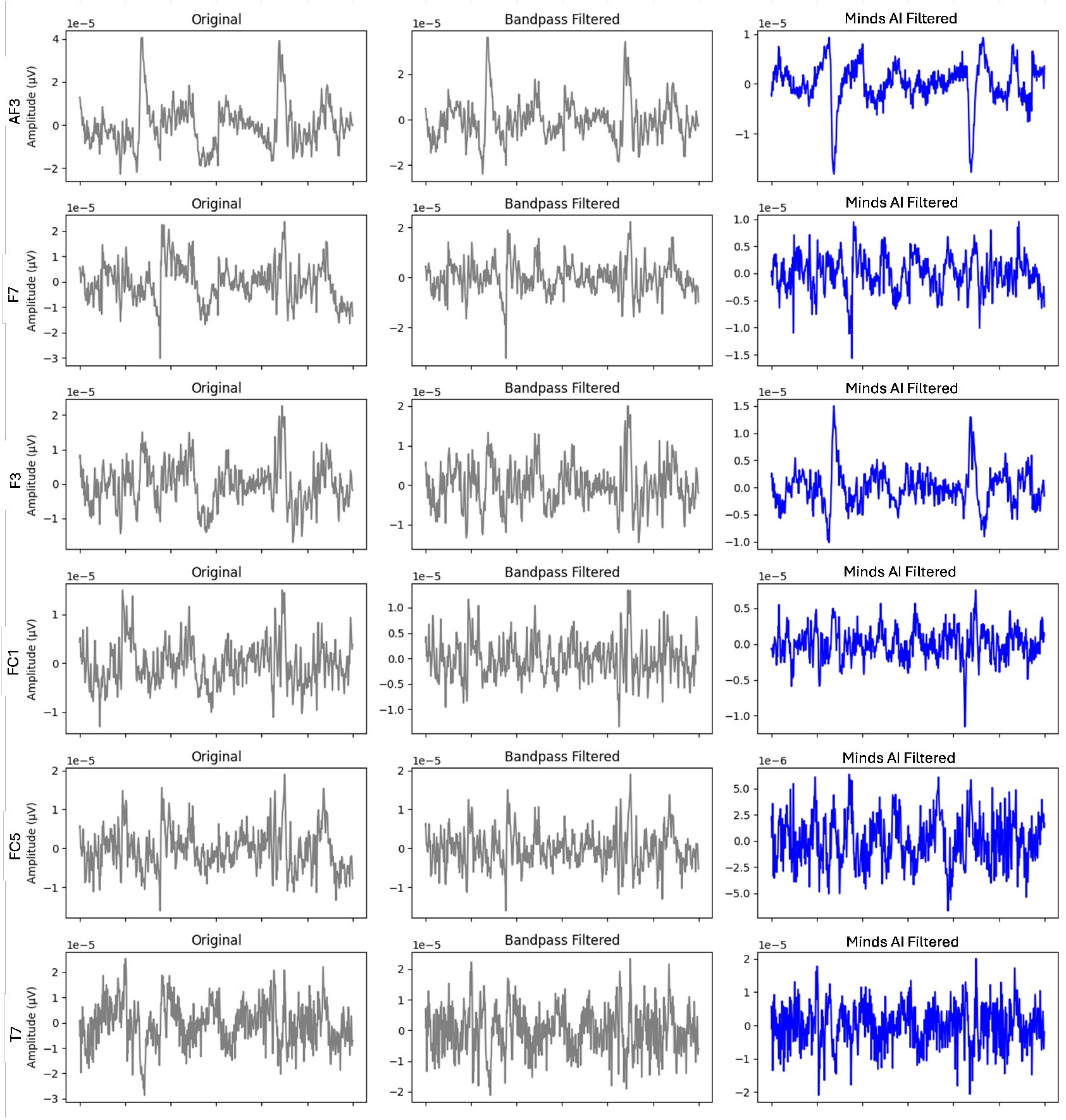
Channel-wise EEG denoising (side-by-side). For each electrode, we show the original signal (left), band-pass baseline (middle), and the Minds AI Filter output (right). The Minds AI Filter suppresses transient motion/ocular bursts while preserving underlying rhythms across channels.

### 5.2 Downstream AI classification (positive vs. negative valence)

We evaluated a two-class valence decoder (positive vs. negative) using 60 s windows. On the Dragon EEG device, the best Minds AI Filter setting achieved participant-wise balanced accuracy up to 0.73 (best *λ* = 10^*−*5^), with typical subject means in the 0.60–0.70 range. Relative to a band-pass–only pipeline, the Minds AI Filter improved balanced accuracy by *∼* 14%–20% on this headset; gains on the separate, prefiltered wet-electrode DEAP dataset were expectedly smaller with an average of (*∼* 6%) but maximum individual gain of (*∼* 35%). We observed no cases where adding the Minds AI Filter reduced downstream classification; when combined with other preprocessing, effects were neutral or positive. Figure 2 summarizes cross-pipeline distributions; subject-wise changes are shown in Fig. 3.

**Figure 2:**
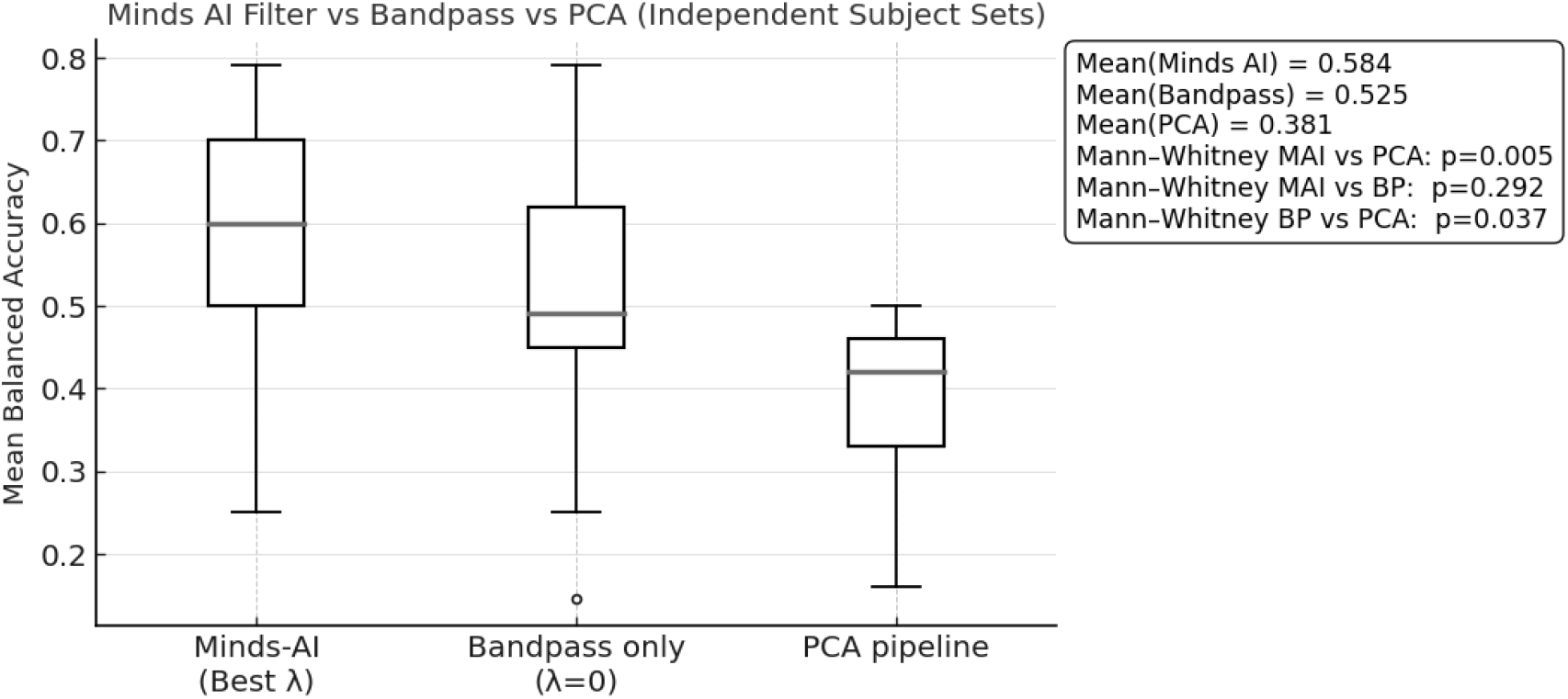
Comparison of mean balanced accuracy across pipelines. Boxes indicate interquartile range with whiskers to non-outlier range; per-pipeline means and Mann–Whitney *p*-values are shown in the inset.

**Figure 3:**
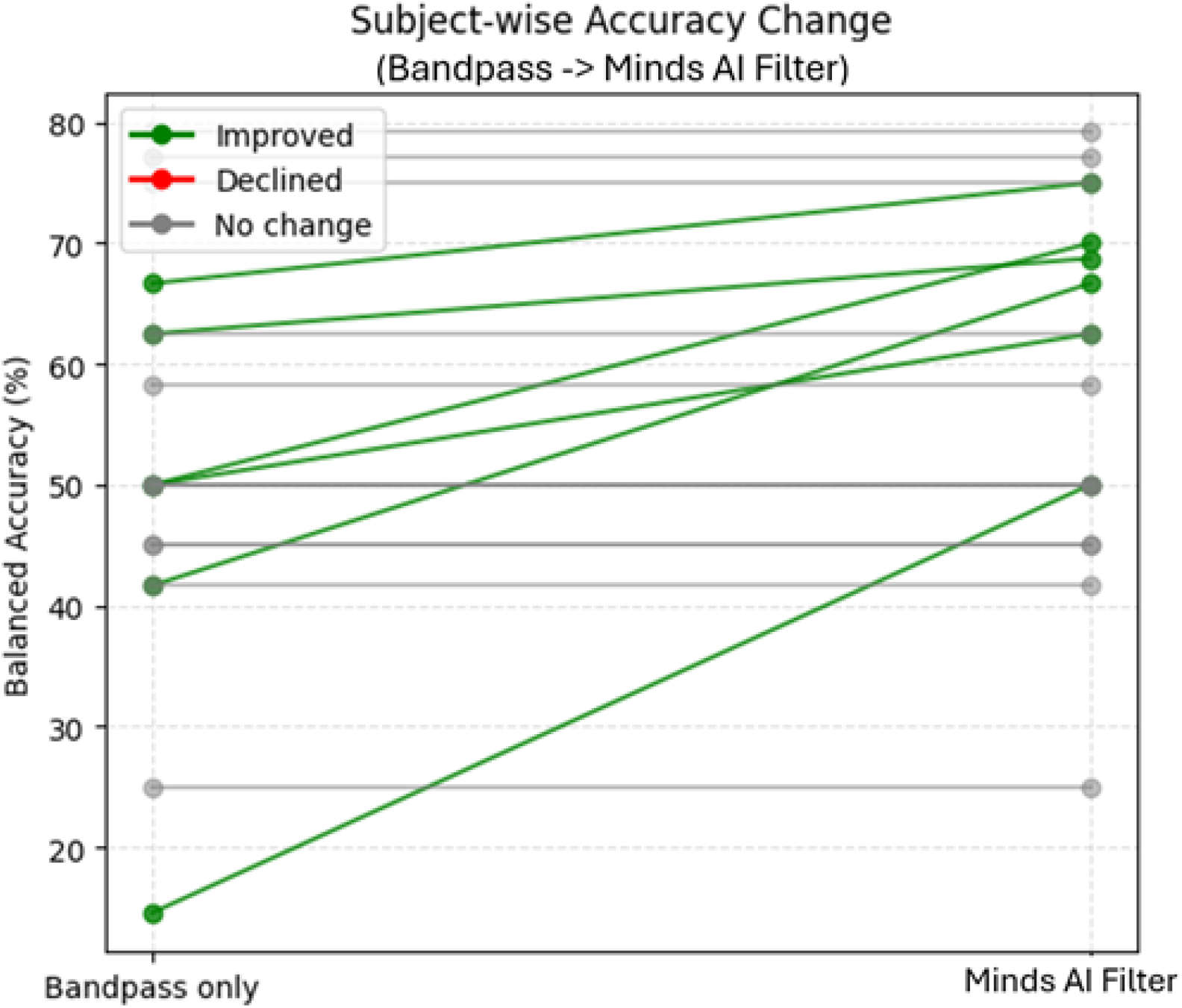
Subject-wise balanced accuracy change from a band-pass–only baseline to the Minds AI Filter–preprocessed pipeline. Each line connects a subject’s performance between conditions; green = improved, red = declined, gray = no change.

### 5.3 Comparisons to established filters

We compared against representative families: (i) **band-pass/notch + CAR/Laplacian** spatial filters; (ii) **PCA/ICA** pipelines with standard component rejection; (iii) **adaptive filters** (e.g., LMS variants) for ocular/motion contamination; and (iv) **ASR-style** artifact subspace methods. In real-time windows, our method maintained higher SNR-like metrics under dynamic artifacts and yielded equal or better downstream classification than each baseline across devices. Offline (longer windows), our method matched or exceeded alternatives while avoiding component-labeling steps. Aggregate comparisons and subject-level distributions are summarized in Figs. 3 and 2.

### 5.4 Latency & real-time viability

Profiling on commodity laptops shows low added latency compatible with online feedback and inference. We report latency as a function of window/hop size and sampling rate, together with throughput and CPU utilization. Per 60 second windows of data: **Mean computational latency:** 0.2637 s **(SD:** 0.1377 s).

## 6 Discussion

The Minds AI Filter provides a practical, device-agnostic improvement for online EEG by leveraging multichannel synchrony with a single hyperparameter. It reduces large, transient artifacts while retaining physiologically plausible structure, which stabilizes downstream features for AI models. Limitations include sensitivity to very sparse montages (*≤*4 electrodes) and the need to tune *λ* with window length and sampling rate. Future work includes adaptive *λ*, data-driven connectivity priors, expanded tasks (motor imagery, inner speech), artifact-specific benchmarks, and validation on other physiological time-series.

## 7 Ethics & Consent

All in-house recordings were obtained with informed consent under protocols compliant with institutional and national guidelines. Details of approvals, recruitment, and anonymization will be provided in the final manuscript.

## 8 Data & Code Availability

We provide **Supplementary File S1** (`filter.txt`) containing a minimal reference implementation sufficient to reproduce the figures/tables, and **Supplementary File S2** (`Minds_AI_Filter_ Real-time_Signal_Analysis.txt`) with the exact real-time metric computations used in this work. The code package and real-time test app (with potential version updates) can be found at https://github.com/MindsApplied/Minds_AI_EEG_Filter. The same files (released by MindsApplied) will be archived with a DOI at submission. Public datasets are cited; in-house data will be shared in de-identified form where permitted.

